# Fishing for mammals: landscape-level monitoring of terrestrial and semi-aquatic communities using eDNA from lotic ecosystems

**DOI:** 10.1101/629758

**Authors:** Naiara Guimarães Sales, Maisie B. McKenzie, Joseph Drake, Lynsey R. Harper, Samuel S. Browett, Ilaria Coscia, Owen S. Wangensteen, Charles Baillie, Emma Bryce, Deborah A. Dawson, Erinma Ochu, Bernd Hänfling, Lori Lawson Handley, Stefano Mariani, Xavier Lambin, Christopher Sutherland, Allan D. McDevitt

## Abstract

1. Environmental DNA (eDNA) metabarcoding has revolutionised biomonitoring in both marine and freshwater ecosystems. However, for semi-aquatic and terrestrial animals, the application of this technique remains relatively untested.
2. We first assess the efficiency of eDNA metabarcoding in detecting semi-aquatic and terrestrial mammals in natural lotic ecosystems in the UK by comparing sequence data recovered from water and sediment samples to the mammalian communities expected from historical data. Secondly, we evaluate the detection efficiency of eDNA samples compared to multiple conventional non-invasive survey methods (latrine surveys and camera trapping) using occupancy modelling.
3. eDNA metabarcoding detected a large proportion of the expected mammalian community within each area. Common species in the areas were detected at the majority of sites. Several key species of conservation concern in the UK were detected by eDNA in areas where authenticated records do not currently exist, but potential false positives were also identified for several non-native species.
4. Water-based eDNA samples provided comparable results to conventional survey methods in per unit of survey effort for three species (water vole, field vole, and red deer) using occupancy models. The comparison between survey ‘effort’ to reach a detection probability of ≥0.95 revealed that 3-6 water replicates would be equivalent to 3-5 latrine surveys and 5-30 weeks of single camera deployment, depending on the species.
5. *Synthesis and Applications*. eDNA metabarcoding represents an extremely promising tool for monitoring mammals, allowing for the detection of multiple species simultaneously, and provides comparable results to widely-used conventional survey methods. eDNA from freshwater systems delivers a ‘terrestrial dividend’ by detecting both semi-aquatic and terrestrial mammalian communities, and provides a basis for future monitoring at a landscape level over larger spatial and temporal scales (i.e. long-term monitoring at national levels).

## Introduction

Environmental DNA (eDNA) metabarcoding (the simultaneous identification of multiple taxa using DNA extracted from an environmental sample, e.g. water, soil, using next-generation sequencing) has revolutionised the way we approach biodiversity monitoring in both marine and freshwater ecosystems (Valentini et al., 2016; Deiner et al., 2017). Successful applications include tracking biological invasions, detecting rare and endangered species and describing entire communities (Deiner et al., 2017). Since water has been shown to be a reliable source of eDNA (Deiner et al., 2017), most eDNA metabarcoding applications to date have been focused on monitoring fishes, amphibians and macroinvertebrates (Fernández et al., 2018; Hänfling et al., 2016; Valentini et al., 2016). What has become apparent from studies in lentic systems (ponds and lakes) is that semi-aquatic and terrestrial mammals can also be detected using universal primer sets for vertebrates, despite not being the focal taxonomic group (Hänfling et al., 2016; Harper et al., 2019). As a result, there has been an increasing focus on the use of both vertebrate (Harper et al., 2019) and mammal-specific primer sets (Ishige et al., 2017; Ushio et al., 2017) for detecting mammalian communities using eDNA metabarcoding.

Mammals include some of the most imperiled taxa, with over one fifth of species considered to be threatened or declining (Visconti et al., 2011), hence identification of *in-situ* biodiversity levels is essential. Given that any optimal survey approach is likely to be species-specific, very few species can be detected at all times when they are present. This imperfect detection (even greater for elusive and rare species) can lead to biased estimates of occurrence and hinder species conservation (Mackenzie et al., 2003). For mammals, repeated surveys using several monitoring methods are usually applied, such as indirect observations of latrines, faeces, hair, or tracks, or direct observations such as live-trapping or camera trapping surveys over short time intervals such that closure/invariance can be assumed and detectability estimated (Nichols et al., 2008). Each of these methods has associated efficiency, cost and required expertise trade-offs, which become more challenging as the spatial and temporal scales increase.

eDNA yields species-specific presence/absence data that are likely to be most valuable for inferring species distributions using well established analytical tools such as occupancy models (MacKenzie et al., 2002). These models resolve concerns around imperfect detection of difficult to observe species and, using location-specific detection histories, can be used to infer true occurrence states, factors that influence occupancy rates, colonization-extinction probabilities, and estimates of detection probability (MacKenzie et al., 2017). The use of eDNA to generate species-specific detection data has unsurprisingly increased in recent years, and in many cases has outperformed or at least matched conventional survey methods (Lugg et al., 2018). Although comparisons between eDNA analysis and conventional surveys for multi-species detection are numerous (see Table S1 in Lugg et al., 2018), studies focusing on detection probability estimates for multiple species identified by metabarcoding are rare (Abrams et al., 2019; Valentini et al., 2016).

The aim of this study was to assess the efficiency of eDNA for detecting semi-aquatic and terrestrial mammals in natural lotic systems in the UK. We conducted eDNA sampling in rivers and streams in two areas (Assynt, Scotland and Peak District National Park, England), which together have the majority of UK semi-aquatic and terrestrial mammalian species present (Table S1). Our objectives were two-fold: first, we sought to establish whether eDNA metabarcoding is a viable technique for monitoring semi-aquatic and terrestrial mammals by comparing it to the mammalian communities expected from historical data, a group for which eDNA sampling has rarely been evaluated in a natural setting. Secondly, we evaluate the detection efficiency of water- and sediment-based eDNA sampling in one of these areas (Assynt) for multiple species compared to multiple conventional non-invasive survey methods (latrine surveys and camera trapping).

## Material and Methods

### Latrine surveys

Assynt, a heather-dominated upland landscape in the far northwest of the Scottish Highlands, UK (Fig. 1A), is the location of an ongoing 20-year metapopulation study of water voles (*Arvicola amphibius*) led by the University of Aberdeen. Here, we focus only on data collected in 2017. The metapopulation is characterized by 116 discrete linear riparian habitat patches (ranging from 90 m to nearly 2.5 km) distributed sparsely (4% of waterway network) throughout the 140 km^2^ study area (Sutherland et al., 2014). Water voles use prominently placed latrines for territory marking (Fig. S1A). Using latrine surveys, a reliable method of detection (Sutherland et al., 2014), water vole occupancy status was determined by the detection of latrines that are used for territory marking (Sutherland et al., 2013). During the breeding season (July and August), latrine surveys were conducted twice at each site. In addition to water vole latrines, field vole (*Microtus agrestis*) pellets are also easily identifiable, and so field vole detections were also recorded along waterways as a formal part of the latrine survey protocol. Live-trapping was then carried out at patches deemed to be occupied by water voles according to latrine surveys to determine their abundances (this was used to determine which sites were sampled for eDNA; see below).

**Figure 1.**
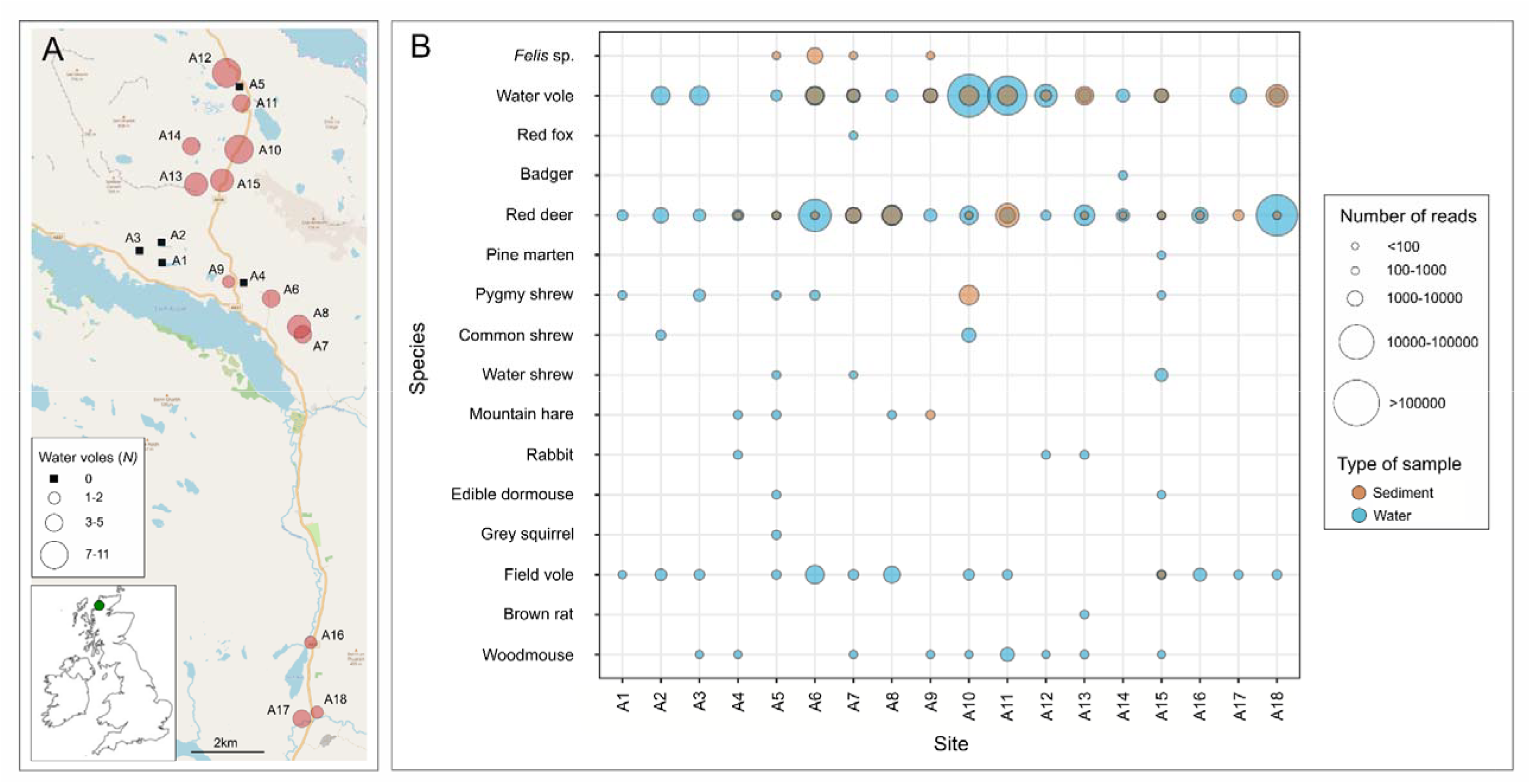
Environmental DNA (eDNA) sampling sites in Assynt, Scotland (A). Categorical values for water vole abundance at each site based on live-trapping data. In (B), a bubble graph representing presence-absence and categorical values of the number of reads retained (after bioinformatic filtering) for eDNA (water in blue and sediment in orange) from each wild mammal identified in each site in Assynt (A1-A18).

### Camera Trap Data

Camera traps were deployed at the beginning of July and thus overlapped temporally with the latrine survey in Assynt. As part of an assessment of the value of cameras for monitoring water voles, data were collected from cameras deployed at seven of these patches for the purpose of this study. Within each of these patches, cameras were deployed at the midpoint of the areas where active signs (latrines, grass clipping, burrows) were detected, and if no signs were detected, at the midpoint of historical water vole activity (J. Drake, C. Sutherland and X. Lambin, *pers. comm*.). These will also capture images of any species present in the area that come within close proximity of the camera (Fig. S2A-F).

Cameras were deployed approximately 1 m above ground on iron ‘u-posts’ to avoid flooding, prevent knock-down by wind/wildlife, and optimize both depth of field and image clarity. Cameras (Bushnell HD Trophy Cam, Bushnell Outdoor Products, Overland Park, Missouri, USA) were set at normal detection sensitivity (to reduce false-triggers from grass/shadows), low night time LED intensity (to prevent image white out in near depth of field), three shot burst (to increase chance of capturing small, fast moving bodies), and 15 min intervals between bursts (to increase temporal independence of captures and decrease memory burden). The area each camera photographed was approximately 1-2 m^2^. Animals were identified on images and information was stored as metadata tags using the R (R Core Team, 2018) package camtrapR following the procedures described in Niedballa et al. (2018). Independence between detections was based on 60-minute intervals between species-specific detections.

### eDNA sampling

A total of 18 potential water vole patches were selected for eDNA sampling in Assynt from 25-27^th^ October 2017. The time lag between the latrine/live-trapping and eDNA surveys was because of two main reasons: (i) legitimate concerns around cross-site DNA contamination during latrine/live-trapping where researchers moved on a daily basis between sites as well as regularly handled and processed live animals and (ii) the selection of eDNA sampling sites was based on the latrine surveys and abundance data provided by live-trapping so could only occur after this was completed by August 6^th^. Water and sediment samples were collected from patches where water voles were determined to be absent (five sites; A1-A5); with 1-2 individuals present (three sites; A9, A16 and 18); 3-5 individuals (five sites; A6, A8, A11, A14 and A17); and 7-11 individuals (five sites; A7, A10, A12, A13 and A15; Fig. 1A). Each of these streams/rivers differed in their characteristics (in terms of width, depth and flow) and a representation of the sites is depicted in Fig. S3A-D. Three water (two litres each) and three sediment (~25mL) replicates were taken at each patch (further details of sample collection are provided in the Supplementary Material: Appendix 1).

In addition to Assynt, eDNA sampling was also conducted on a smaller scale in the Peak District National Park, England (Fig. S4) to incorporate additional mammals that are not known to be present in Assynt (Table S1). Here, the occurrence of water vole was identified by the presence of latrines in two sites (P1 and P2) at the time of eDNA sampling (Fig. S1A), whilst no latrines were identified at one site (P3). At site P1, an otter (*Lutra lutra*) spraint was identified at the time of eDNA sampling (Fig. S1B). These three sites were sampled in March 2018 using the same methodology as in Assynt but were taken in close proximity (<50cm) to water vole latrines where present (Fig. S1A).

### eDNA Laboratory Methods

DNA was extracted from the sediment samples using the DNeasy PowerMax Soil kit and from the water samples using the DNeasy PowerWater Kit (both QIAGEN Ltd.) in a dedicated eDNA laboratory in the University of Salford. In order to avoid the risk of contamination during this step, DNA extraction was conducted in increasing order of expected abundance in the eDNA samples (all field blanks were extracted first, followed by the sites with supposedly zero water vole abundance, up to the highest densities last). Along with field blanks (Assynt = 8, Peak District = 2), six lab extraction blanks were included (one for each daily block of extractions). A decontamination stage using a Phileas 25 Airborne Disinfection Unit (Devea SAS) was undertaken before processing samples from different locations. eDNA was amplified using the MiMammal 12S primer set (MiMammal-U-F, 5′-GGGTTGGTAAATTTCGTGCCAGC-3′; MiMammal-U-R, 5′-CATAGTGGGGTATCTAATCCCAGTTTG-3′) (Ushio et al., 2017) targeting a ∽170bp amplicon from a variable region of the 12S rRNA mitochondrial gene. A total of 147 samples, including field collection blanks (10) and laboratory negative controls (12, including six DNA extractions blanks and six PCR negative controls), were sequenced in two multiplexed Illumina MiSeq runs. Details of PCR conditions, library preparation and bioinformatic analyses are provided in Appendix 1 in the Supplementary Material.

### Occupancy/Detection Analysis in Assynt

The data collection from the different survey types described above (water-based eDNA, sediment-based eDNA, latrine and camera traps) produced the following site-specific detection/non-detection data:

a. Latrine: two latrine surveys at 116 patches.
b. w-eDNA: three water-based eDNA samples at 18 of the 116 patches surveyed.
c. s-eDNA: three sediment-based eDNA samples at 18 of the 116 patches surveyed.
d. Camera: six one-week occasions of camera trapping data at seven of the 18 patches surveyed by both Latrine and eDNA (w-eDNA + s-eDNA) surveys.

We chose to focus on three species that were detected by at least three of the four methods: water voles, field voles and red deer (*Cervus elaphus*). Water voles and field voles were recorded using all four survey methods and had detection histories for 14 surveying events ((Latrine × 2) + (w-eDNA × 3) + (s-eDNA × 3) + (Camera × 6)). Red deer were not recorded during latrine surveys and had detection histories for 12 surveying events ((w-eDNA × 3) + (s-eDNA × 3) + (Camera × 6)). To demonstrate the relative efficacy of the four surveying methods, we restricted the analyses to the 18 sites where both latrine surveys were conducted and eDNA samples were taken, seven of which had associated camera trapping data. Although each surveying method differs in terms of effort and effective area surveyed, each are viable surveying methods that are readily applied in practice. So, while the specific units of effort are not directly comparable, the relative detection efficacy per surveying method-specific unit of effort is of interest and will provide important context for designing future monitoring studies and understanding the relative merits of each surveying method. Analyzing the data using occupancy models allowing for method-specific detectability enables such a comparison in per unit effort efficacy between eDNA metabarcoding and multiple conventional survey methods.

A single season occupancy model (MacKenzie et al., 2002) was applied to the ensemble data where detection histories were constructed using each of the surveying events as sampling occasions (MacKenzie et al., 2017). The core assumption here is that the underlying occupancy state (i.e. occupied or empty) is constant over the sampling period, and therefore, every sampling occasion is a potentially imperfect observation of the true occupancy status. Because occasions represent method-specific surveying events, we used “surveying method” as an occasion-specific covariate on detection (survey method type: Latrine, w-eDNA, s-eDNA and Camera). Our primary objective was to quantify and compare method-specific detectability, so we did not consider any other competing models. For comparing the methods, we compute accumulation curves as (MacKenzie & Royle, 2005):

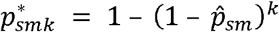

Where 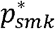 is the cumulative probability of detecting species *s*, when species *s* is present, using method *m* after *k* surveying events based on the estimated surveying method-specific detection probability for each species 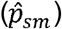. We vary k from 1 to a large number and find the value of *k* that results 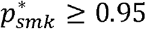. We conducted the same analysis separately for water voles, field voles, and red deer. Analysis was conducted in R (R Core Team, 2018) using the package unmarked (Fiske & Chandler, 2011).

## Results

### Mammal Detection via eDNA metabarcoding

A total of 125 eDNA samples and 22 field and laboratory controls were sequenced. The two sequencing runs generated 23,276,596 raw sequence reads and a total of 15,463,404 sequences remained following trimming, merging, and length filtering. Vertebrate species that were likely contaminants from other projects on South American freshwater and European marine-based projects in the lab were excluded. After bioinformatic analysis, the final ‘filtered’ dataset contained 23 mammals (Tables S2 and S3).

For mammals, ~12 million reads were retained after applying all quality filtering steps (see Appendix 1). Reads from humans represented, along with cattle (*Bos taurus*), pig (*Sus scrofa*), horse (*Equus ferus*), sheep (*Ovis aries*) and dog (*Canis lupus familiaris*), were not considered further as the focus of this study was on wild mammals (Table S4). *Felis* was included because of the potential of it being wildcat (*Felis silvestris*) or domestic cat (*F. catus*)/wildcat hybrids. A final dataset comprising ~5.9 million reads was used for the downstream analyses (Table S4).

In Assynt, the wild species identified were the red deer (18/18 sites); water vole (15/18); field vole (13/18); wood mouse (*Apodemus sylvaticus* - 9/18); pygmy shrew (*Sorex minutus* - 4/18); wild/domestic cat (*Felis* spp. - 4/18); mountain hare (*Lepus timidus* - 4/18); rabbit (*Oryctolagus cuniculus* - 3/18); water shrew (*Neomys fodiens* - 3/18); common shrew (*Sorex araneus* - 2/18); edible dormouse (*Glis glis* - 2/18); grey squirrel (*Sciurus carolinensis* - 1/18); pine marten (*Martes martes* - 1/18); brown rat (*Rattus norvegicus* - 1/18); red fox (*Vulpes vulpes* - 1/18) and badger (*Meles meles* - 1/18; Fig. 1B). All of these species are distributed around/within Assynt (Table S1), with the exception of the edible dormouse and the grey squirrel. These are unequivocally absent from the region. The edible dormouse is only present in southern England and the grey squirrel is not distributed that far north in Scotland (Mathews et al., 2018).

Of the wild mammals in the Peak District, the water vole, field vole, wood mouse and otter were found in two sites (P1 and P2). The red deer, pygmy shrew, common shrew, water shrew, red squirrel (*Sciurus vulgaris*), grey squirrel, pine marten and badger were each found at a single site (Fig. S4). Only the rabbit was found in site P3, the site chosen on the expectation that water voles and otters would not be found (S. Proctor, *pers. comm*.). All species identified are currently distributed within the Park (Table S1), except the red squirrel and pine marten. The pine marten, which is critically endangered in England, has only two reliable records that have been confirmed in the Park since 2000 and the red squirrel has not been present for over 18 years (Alston et al. 2012).

Overall, water samples yielded better results than sediment samples regarding species detection and read count for both areas sampled (Figs 1B and S4). In Assynt, only the wild/domestic cat was exclusively detected in sediment samples (four sites), whereas water samples recovered eDNA for ten additional species not found in the sediment samples. The red deer, water vole, field vole, mountain hare and pygmy shrew were also found in sediment samples in Assynt (Fig. 1B), and water vole and wood mouse in the Peak District sediment samples (Fig. S4).

### Occupancy Analysis

Of the 18 sites where both latrine and eDNA surveys were conducted, water voles were detected at 13, and field voles were detected at 11. A total of seven wild mammals were recorded at the seven sites with a camera trap from July 10^th^ to October 25^th^ 2017 (Fig. S2 and Table S5). There were several incidences where a shrew could not be identified to species level using camera traps. For camera traps, water voles were recorded at all sites, red deer at five out of seven, field voles and weasels at three sites, water shrews and otters at two, and a red fox at a single site.

For the 18 sites in Assynt, estimated site occupancy (with 95% confidence intervals) from the combined surveying methods was 0.91 (0.63 – 0.98) for water voles and 0.88 (0.57 – 0.98) for field voles. Red deer were observed at every patch by at least one of the methods, and therefore occupancy was 1 (Table 1). For all three species, per sample detection probability was higher for eDNA taken from water than for eDNA taken from sediment (Table 1, Fig. 2). The surveying method specific efficacy pattern was similar for water voles and field voles (Table 1, Fig. 2): latrine surveys had the highest probability of detecting the species (0.77 and 0.52 respectively), followed by eDNA from water (0.57 and 0.40 respectively), then camera trapping (0.50 and 0.20 respectively), and finally eDNA from sediment (0.27 and 0.02 respectively). Detection probability was higher for water voles than field voles using all four methods (Table 1, Fig. 2). No effort was made to record red deer presence during latrine surveys. Like the water voles and field voles, red deer detection has higher using eDNA from water (0.67, CI: 0.53 – 0.78) compared to eDNA from sediment (0.10, CI: 0.04 – 0.21). Unlike the voles, which were more detectable by cameras than sediment eDNA, red deer detection on cameras was similar to sediment eDNA (0.10, CI: 0.04 – 0.24).

**Table 1.** Estimated site occupancies and detection probabilities obtained for water-based eDNA (w-eDNA), sediment-based eDNA (s-eDNA) and conventional survey methods (Latrine and Camera) in Assynt.

| Species | Occupancy | Detection probability |  |  |  |
| --- | --- | --- | --- | --- | --- |
|  |  | *Latrine* | *w-eDNA* | *s-eDNA* | *Camera* |
| Water vole | 0.91<br>(0.63 – 0.98) | 0.77<br>(0.59 – 0.89) | 0.57<br>(0.43 – 0.71) | 0.27<br>(0.16 – 0.41) | 0.50<br>(0.35 – 0.65) |
| Field vole | 0.89<br>(0.57 – 0.98) | 0.52<br>(0.34 – 0.69) | 0.40<br>(0.26 – 0.55) | 0.02<br>(0.00 – 0.14) | 0.20<br>(0.10 – 0.37) |
| Red deer | 1.00<br>(1.00 – 1.00) | -- | 0.67<br>(0.53 – 0.78) | 0.10<br>(0.04 – 0.21) | 0.10<br>(0.09 – 0.24) |

**Figure 2.**
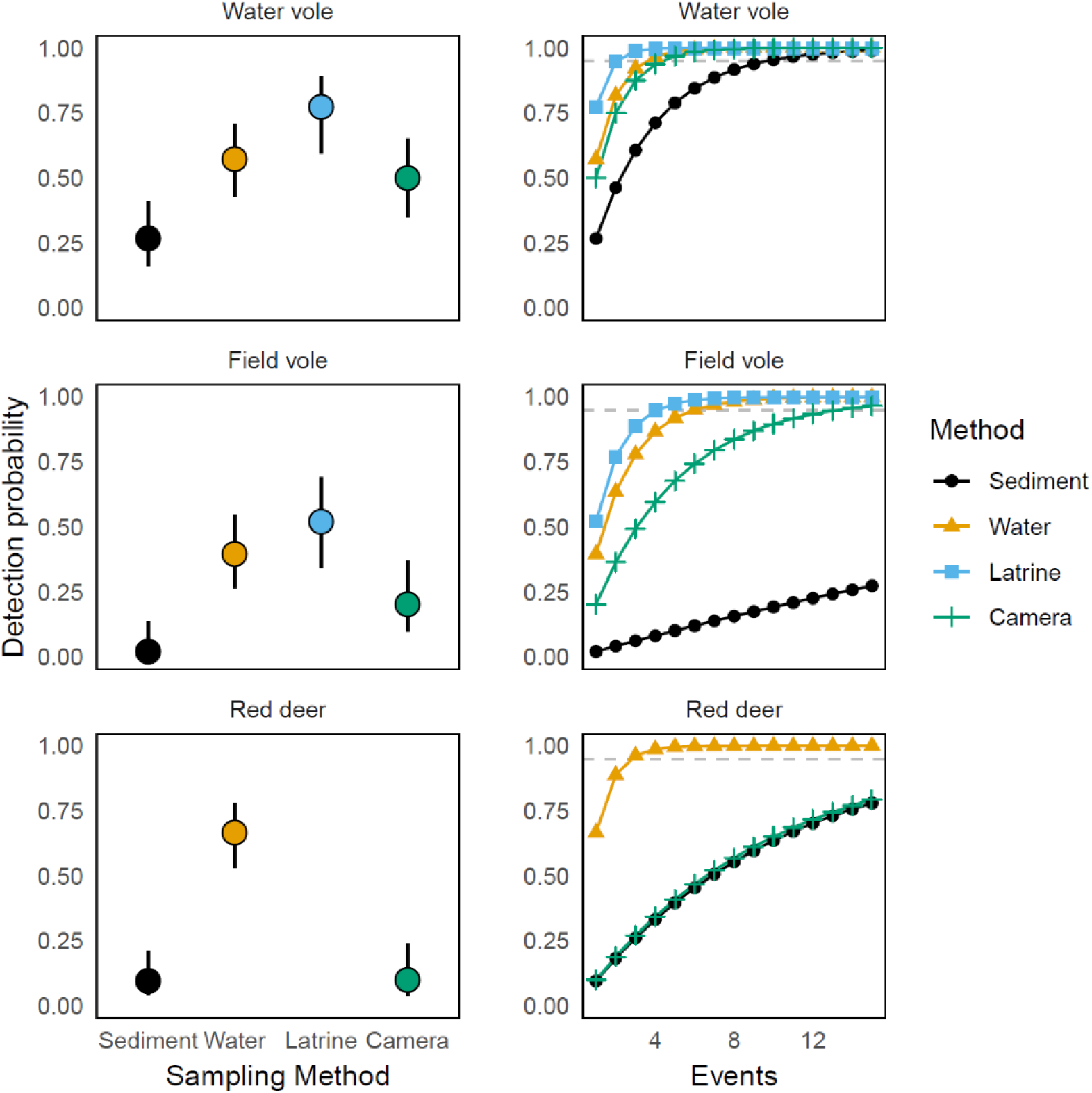
The detection probabilities of each survey method (sediment-based eDNA, water-based eDNA, latrine and camera) for each of three focal species (from top to bottom on the left); water vole; field vole and red deer. On the right, the accumulation curves for each species for the number of sampling events for each survey method to provide a ≥0.95 probability of detection.

The patterns described above detail surveying event-specific detectability. We also computed the cumulative detection probability for each method and each species 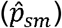. The cumulative detection curves over 15 surveying events are shown in Fig. 2. The number of surveying events, *k*, required to achieve 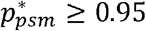 for water voles was 3 surveys, 4 samples, 10 samples, and 5 weeks, for latrines, water eDNA, sediment eDNA, and cameras respectively. The number of surveying events, *k*, required to achieve 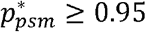 for field voles was 5 surveys, 6 samples, 141 samples, and 14 weeks, for latrines, water eDNA, sediment eDNA, and cameras respectively. The number of surveying events, *k*, required to achieve 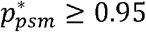 for red deer was 3 samples, 30 samples, and 29 weeks, for water eDNA, sediment eDNA, and cameras respectively (see also Fig. 2).

## Discussion

Despite the increasing potential of eDNA metabarcoding as a biomonitoring tool (Deiner et al., 2017), its application has largely been focused on strictly aquatic or semi-aquatic animals, thus restricting management and conservation efforts of the wider ecosystem (Williams et al., 2018). Here, we demonstrate the ability of eDNA metabarcoding to provide a valuable ‘terrestrial dividend’, mammals in this case, from a freshwater lotic system, with a large proportion of the expected species from the wider ecosystem being detected in each of the two study locations. In particular, we have demonstrated that water-based eDNA offers a promising and complementary tool to conventional survey methods for the detection of whole mammalian communities.

### Detection of mammalian communities using eDNA metabarcoding

Of the species known to be common in both Assynt and the Peak District, eDNA metabarcoding readily detected the water vole, field vole and red deer at the majority of sites surveyed (Figs. 1B and S4). The pygmy, common and water shrews, wood mice and mountain hares were also detected by eDNA metabarcoding at multiple sites in Assynt (Fig. 1B). A higher eDNA detection rate is expected for aquatic and semi-aquatic mammals compared to terrestrial mammals in aquatic environments due to the spatial and temporal stochasticity of opportunities for terrestrial mammals to be in contact with the water (Ushio et al., 2017). The semi-aquatic water vole was generally detected by eDNA metabarcoding where we expected to find it and at relatively high read numbers (Figs. 1B and S4), in line with previous studies in lentic systems (Harper et al., 2019). However, the red deer was the only terrestrial species detected by eDNA at all sites in Assynt, and the terrestrial field vole at over 70% of surveyed sites.

In addition to lifestyle (semi-aquatic or terrestrial), the number of individuals of each species (i.e. group-living) may be important for eDNA detection (Williams et al., 2018). As a counter example to this, otters and weasels were notably absent in the eDNA samples in Assynt despite being picked up by camera traps (Fig. S2 and Table S5). Otters were present in the water eDNA samples at two sites in the Peak District, albeit at a lower number of reads in comparison to most of the other species detected (Fig. S4; Table S2). This mirrors previous studies where eDNA analysis has performed relatively poorly for otter detection in captivity and the wild (Harper et al., 2019; Thomsen et al., 2012). Carnivores were generally detected on fewer occasions (e.g. red foxes, badgers and pine martens; Figs. 1B and S4) or not at all (e.g. stoats and American mink in addition to those discussed above) in comparison to smaller mammals and red deer. For some of these species, a relatively large home range and more solitary nature (e.g. red foxes) may go some way towards explaining a lack of, or few, eDNA records.

Regarding the sampling medium for eDNA, here we demonstrated that water is a more effective method for detection of mammal eDNA than sediment (Table 1; Figs. 1B and S4). For one of our focal species, the water vole, 75% of sites which were deemed unoccupied by latrine surveys and those with ≤2 individuals (8 sites) in Assynt, returned a non-detection for sediment eDNA as opposed to 37.5% of sites for water (Figs. 1A and 1B). It is worth investigating further if sediment eDNA could indicate the presence of a more ‘established’ population, where a certain threshold of individuals and long-term occupation (i.e. historical) is required for detection in sediment (Turner et al., 2015).

Importantly, sparse or single eDNA records can be significant. The edible dormouse and grey squirrel sequences identified within the Assynt samples (Fig. 1B) and red squirrel within the Peak District (Fig. S4) highlights the caveats associated with this technique. Should management have relied on eDNA evidence alone, as the edible dormouse and grey squirrel are classified as invasive species within Great Britain, false positives for these species could lead to unnecessary resources being allocated for management/eradication programmes. These likely arose due to cross-contamination between labs from reference database construction. Controlling for false positives is certainly a huge challenge in eDNA metabarcoding (Ficetola et al., 2015). Even with these concerns around false positives highlighted, two records are potentially noteworthy in a conservation context for UK mammals because of the relatively high read number associated with these records (Tables S2 and S3). The first of these is the *Felis* records in sediment samples in multiple sites in Assynt (Fig. 1B). Even with ‘pure’ *F. silvestris* as reference sequences, it was not possible to distinguish between the wild and domesticated species for this 12S fragment (data not shown). Despite ongoing conservation efforts, there may now be no ‘pure’ Scottish wildcats left in the wild in the UK (Senn et al., 2018) but isolated populations (perhaps of hybrid origin) may exist in this region (Sainsbury et al., 2019). The other significant eDNA record was the pine marten in the Peak District. The pine marten had disappeared from most of the UK but has been recovering from historical persecution. However, authentic records from northern England are scarce or lacking altogether (Alston et al., 2012; Sainsbury et al., 2019). The pine marten is a threatened in the UK but is potentially expanding its range. There is record of a recent roadkill exists from just outside the Park’s boundary (BBC News, 2018).

### Comparisons between surveying methods

Comparisons of species detection by traditional survey approaches and eDNA analysis are now numerous in the literature, and mainly focus on what is and what is not detected within and across different methods (Hänfling et al., 2016; Lugg et al., 2018). Yet, there has been growing incorporation of occupancy modelling to estimate the probability of detecting the focal species, in comparison to one other survey method, either for a single species (Lugg et al., 2018) or multiple species (Valentini, et al., 2016; Abrams et al., 2019). Simultaneous multi-method comparisons for multiple species have been lacking and this study directly addresses this for the first time.

The probability of detecting the water vole and field vole was higher for the latrine surveys than eDNA sampling (both water and sediment) and camera traps (Table 1; Fig. 2). However, when considering confidence intervals, there was considerable overlap between latrine, water-based eDNA and camera traps for both species, with only sediment-based eDNA yielding a low probability of detection (Table 1). Detection probabilities for water-based eDNA and camera traps were similar for water voles, with camera traps less likely to detect the field vole than water-based eDNA. For the red deer (for which no latrine survey was undertaken), water-based eDNA had a much higher probability of detection than either sediment-based eDNA or camera traps (which performed similarly; Table 1). Despite the increasing adoption of camera traps in providing non-invasive detections for mammals (Hofmeester et al., 2019), camera traps were outperformed by water-based eDNA metabarcoding for the three focal species in this component of the study. Camera traps are certainly limited by their photographic range and placement (amongst many other factors; Hofmeester et al., 2019). Here, camera traps were deployed so as to sample the habitat of the water vole (see Fig. S2), which may explain lower detection for other terrestrial species in comparison to eDNA metabarcoding. It is also worth repeating that eDNA metabarcoding did not detect the otter and weasel which were detected by camera traps and are ubiquitous in the Assynt area (Table S5; Fig. S2). This is similar to the findings of Harper et al., (2019) with the red fox and badger near ponds. Studies focusing on a single species often report that eDNA analysis outperforms the conventional survey method in terms of detection probabilities (e.g. Lugg et al., 2018). Multi-species metabarcoding studies may trade-off a slightly lower (but comparable) detection probability than other survey methods for individual species (Fig. 2) in favour of a better overall “snapshot” of occupancy of the whole mammalian community (Ushio et al., 2017).

The comparison between survey ‘effort’ for the four methods to reach a probability of detection of ≥0.95 is highly informative and provides a blueprint for future studies on mammal monitoring. For the water vole, three latrine surveys would be required. A total of four water-based and 10 sediment-based eDNA replicates or five weeks of camera trapping would be required to achieve the same result (Fig. 2). This increases for the field vole in the same habitat, with five latrine surveys and six water-based eDNA replicates. Sediment-based eDNA would be impractical for this species and camera trapping would take 14 weeks. The red deer would require three water-based eDNA replicates and 29-30 events for sediment-based eDNA and camera trap detection.

What is important here is the spatial component and the amount of effort involved in the field. Taking 4-6 water-based eDNA replicates from around one location within a patch could provide the same probability of detecting these small mammals with three latrine surveys. In many river catchments, there may be 100s to 1000s of kilometres to survey that would represent suitable habitat, and only a fraction of that may be occupied by any given species. This is particularly relevant in the context of recovery of water vole populations post-translocation or in situations where remnant populations are bouncing back after invasive American mink (*Neovison vison*) control has been instigated. On a local scale, finding signs of water voles through latrine surveys is not necessarily difficult, but monitoring the amount of potential habitat (especially lowland) for a species which has undergone such a massive decline nationally is a huge undertaking (Morgan et al., 2019). The use of eDNA metabarcoding from freshwater systems to generate an initial, coarse and rapid ‘distribution map’ for vertebrate biodiversity (and at a relatively low cost) could transform biomonitoring at a landscape level. Then, on the basis of this, practitioners could zoom in to further investigate specific areas for confirmation of rare or invasive species for example.

It is clear that eDNA metabarcoding is a promising tool for monitoring semi-aquatic and terrestrial mammals in both lotic (this study) and lentic systems (Harper et al., 2019; Ushio et al., 2017). We detected a large proportion of the expected mammalian community (Table S1), including the possible presence of priority species. Water-based eDNA is comparable or out-performs other non-invasive survey methods for several species (Fig. 2). However, there remain challenges for the application of this technique over larger spatial and temporal scales. Technical issues of metabarcoding in laboratory and bioinformatic contexts have been dealt with elsewhere (Harper et al., 2019) but understanding the distribution of eDNA transport in the landscape and its entry into natural lotic systems is at an early stage. The characteristics of streams and rivers undoubtedly influence eDNA transportation through the environment (Pont et al., 2018). This clearly requires more detailed and systematic eDNA sampling than undertaken here, particularly in an interconnected river/stream network with organisms moving between aquatic and terrestrial environments. Nonetheless, with a deeper understanding of these mechanisms and species ecologies, it could be feasible that sampling a few key areas (e.g. larger rivers and lakes) within a catchment area could provide data on a large proportion (if not all) of the species within it, even when some species are present at low densities (Deiner et al., 2017). In this regard, future studies might also investigate the value of citizen science, where trained volunteers can contribute to data collection at key sites, thus scaling the reach of research whilst raising public awareness and significance of mammalian conservation concerns though public participation in scientific research (Parsons et al., 2018).

## Data accessibility

Data will be made available on a public repository upon acceptance.

## Authors contributions

ADM, XL, CS, OSW, IC, SM, NGS, EO, BH and LLH conceived the study. Monitoring and live-trapping of water voles was part of XL, CS and JD’s ongoing work on water vole metapopulation dynamics in Assynt. JD and EB carried out the latrine surveys and live-trapping. JD analysed the camera trap data in Assynt. DAD provided information and data on mammals in the Peak District. ADM, NGS, SSB and MBM carried out the eDNA sampling. MBM, NGS, SSB, CB and ADM performed the laboratory work. NGS, OSW, LRH, MBM, CB and ADM carried out the bioinformatic analyses. ADM, NGS, IC and MBM analysed the eDNA data. CS and JD conducted the occupancy modelling. ADM, NGS, CS, JD, MBM and LRH wrote the paper, with all other co-authors contributing to editing and discussions.

## Acknowledgements

The eDNA component of this project was funded by a Small Research Grant from the British Ecological Society (grant no. SR17/1214) and a University of Salford Internal Research Award awarded to ADM. JD was supported by a University of Massachusetts Organismal and Evolutionary Biology Research Grant, University of Massachusetts Spring 2018 Graduate School Fieldwork Grant). We thank Kristy Deiner for enlightening conversations about these results. We are grateful to Jerry Herman and Andrew Kitchener for organizing and preparing the tissue samples from National Museums Scotland. Christine Gregory and Sarah Proctor kindly provided water vole and otter information for sampling in the Peak District. We thank the various landowners for permission to sample on their property.

