## Supplementary Material for "Fishing for mammals: landscape-level monitoring of terrestrial and semi-aquatic communities using eDNA from lotic ecosystems"

**Appendix 1**

**eDNA sample collection**

Three water sample replicates (two litres each) and three sediment sample replicates (50 ml falcon tube, approximately half-filled) were taken at each site in Assynt, always within a reachable distance from the river’s edge and at a depth where sediment samples could be taken (Fig. S3A). Water samples were filtered on site using a Sterivex 0.45 µm filter unit (Merck Millipore) and filters were stored in silica beads in the field (1-3 days) (Majaneva et al., 2018) then frozen until DNA extraction. Sediment samples were stored in 100% ethanol. Appropriate decontamination precautions were taken (use of disposable gloves and all equipment and surfaces were decontaminated by using 50% bleach solution) and collection, extraction and PCR negative controls were included. Samples from the Peak District were filtered within 5 hours in the University of Salford laboratory facilities due to its close proximity to the sampling locations. A single filter was used for each replicate in Assynt and the Peak District, and the volume filtered varied between each, ranging from 150 ml to 2 L (see Tables S2 and S3). The amount of sediment collected also varied, with 4 to 10g used in the extractions. A Pearson’s correlation was performed to determine if the amount of water/sediment influenced the amount of retained reads for mammals after bioinformatic filtering.

**Reference database**

Given that this project proposed to use mammal-specific primers (MiMammal-U, Ushio et al., 2017) to target the same region of 12S as the MiFish primers, an *in silico* evaluation was first performed using ecoPCR (Ficetola et al., 2010) against a custom, phylogenetically curated reference database for mammals distributed in the UK and Ireland. This database was one of several databases constructed for UK vertebrates and used in an eDNA metabarcoding study of pond biodiversity (see Harper et al. 2019 for details). The mammal database was updated in July 2018 for the purposes of the present study. Parameters were set to allow a fragment size of 50-250 bp and different number of mismatches (0, 1, 2, 3) between each primer and each sequence in the reference database. Reference sequence data was available for 103 mammal species (91.96%) in the UK. The nine species that were not represented were either cetaceans or bats. Of those species with reference sequence data (N = 103), 44 (42.72%), 65 (63.11%), 72 (69.90%), and 82 (79.61%) mammals were amplified when 0, 1, 2, and 3 primer-sequence mismatches were allowed respectively. Species that did not amplify under any scenario and of relevance to this study were the European water vole (*Arvicola amphibius*), greater white-toothed shrew (*Crocidura russula*), Millet’s shrew (*Sorex coronatus*), Eurasian pygmy shrew (*Sorex minutus*), field vole (*Microtus agrestis*), common vole (*Microtus arvalis*), grey squirrel (*Sciurus carolinensis*), and European polecat (*Mustela furo*).

Targeting a fragment of the 12S gene, a reference database of 32 UK terrestrial mammals was created from ethanol-preserved tissues samples obtained from National Museums Scotland (Table S6). DNA was extracted using the ISOLATE II kit according to the manufacturer’s protocol. These DNA samples were then included in a large barcoding project using the MiFish (Miya et al., 2015) primers (O. Wangensteen et al., *unpublished data*). Of these mammals, only *Sorex araneus* and *Neomys fodiens* failed to generate sequences. PCRs were then carried out on a subset of the tissue-extracted DNA (see Table S6) and Sanger-sequenced (Macrogen Inc.). Amplicons of 172bp from a variable region of the mitochondrial 12S rRNA gene were obtained with the MiMammal-U primers (Ushio et al., 2017).

***eDNA Laboratory Methods***

A set of 96 primers pairs with seven-base sample-specific oligo-tags and a variable number (2-4) of fully degenerate positions (leading Ns) to increase variability in amplicon sequences were used. PCR amplification was conducted using a single-step protocol and to minimize bias in individual reactions, PCRs were replicated three times for each sample and subsequently pooled. The PCR reaction consisted of a total volume of 20 µl including 10 µl Amplitaq; 0.16 µl of BSA; 1.0 µl of each of the two primers (5 µM); 5.84 µl of ultra-pure water, and 2 µl of DNA template. The PCR profile included an initial denaturing step of 95°C for 10 min, 40 cycles of 95°C for 30s, 60°C for 45s, and 72°C for 30s and a final extension step of 72°C for 5 min. Amplification were checked through electrophoresis in a 1.5% agarose gel stained with GelRed (Cambridge Bioscience). PCR products were pooled in two different sets and a left-sided size selection was performed using 1.1x Agencourt AMPure XP (Beckman Coulter). Illumina libraries were built from each set, using a NextFlex PCR-free library preparation kit according to the manufacturer’s protocols (Bioo Scientific). Libraries were then quantified by qPCR using a NEBNext qPCR quantification kit (New England Biolabs) and pooled in equimolar concentrations along with 1% PhiX (v3, Illumina). The libraries were run at a final molarity of 9pM on an Illumina MiSeq platform in a single MiSeq flow cell using the 2 x 150bp v2 chemistry.

***Bioinformatic analysis***

OBITools metabarcoding package (Boyer et al., 2016) was used for the bioinformatic analysis. Quality of the reads was assessed using FastQC, paired-end reads were aligned using illuminapairedend and the ngsfilter command was used for dataset demultiplexing. Short fragments originated from library preparation artefacts (primer-dimer, non-specific amplifications) and reads containing ambiguous bases were removed applying a length filter selecting fragments of 140-190bp using obrigrep. Clustering of strictly identical sequences was performed using obiuniq and a chimera removal step was applied in vsearch (Rognes et al., 2016) through the uchime-denovo algorithm (Edgar et al., 2011). The taxonomic assignment was conducted using ecotag.

A stringent approach was applied to our analyses to avoid false positives and exclude MOTUs/reads putatively belonging to sequencing errors or contamination. The final dataset included only MOTUs that could be identified to species level (>0.98), and MOTUs containing less than 10 reads and with a similarity to a sequence in the reference database lower than 98% were discarded (Cilleros et al., 2018). Singleton reads within individual replicates were also discarded. The maximum number of reads detected in the controls for each MOTU in each sequencing run were removed from all samples (Table S7). For water voles, field voles and red deer (the most abundant wild mammals in terms of sequence reads in our dataset), this equated to a sequence frequency threshold of ≤0.17%, within the bounds of previous studies on removing sequences to account for contamination and tag jumping (Cilleros et al., 2018; Schnell, Bohmann, & Gilbert, 2015). The number of retained reads per replicate was not significantly correlated with the volume of water filtered (Pearson’s correlation: *r* = 0.213; *p* = 0.094) or the amount of sediment collected (Pearson’s correlation: *r* = 0.076; *p* = 0.556).

**TABLES**

**Table S1.** Species (and the Order to which they belong) that are expected to be found within Assynt (based on Matthews et al. 2018 and the Peak District (Alston et al. 2012) and whether or not they were detected by eDNA. A * indicates species where presence is uncertain from Matthews et al. (2018).

| **Common name** | **Scientific name** | **Order** | **eDNA** |
| --- | --- | --- | --- |
| **Assynt** |  |  |  |
| Red deer | *Cervus elaphus* | *Artiodactyla* | **Yes** |
| Sika deer | *Cervus nippon* | *Artiodactyla* | No |
| Roe deer | *Capreolus capreolus* | *Artiodactyla* | No |
| Water vole | *Arvicola amphibius* | *Rodentia* | **Yes** |
| Field vole | *Microtus agrestis* | *Rodentia* | **Yes** |
| Wood mouse | *Apodemus sylvaticus* | *Rodentia* | **Yes** |
| Bank vole* | *Myodes glareolus* | *Rodentia* | No |
| Brown rat | *Rattus norvegicus* | *Rodentia* | **Yes** |
| Pygmy shrew | *Sorex minutus* | *Eulipotyphla* | **Yes** |
| Water shrew | *Neomys fodiens* | *Eulipotyphla* | **Yes** |
| Common shrew | *Sorex araneus* | *Eulipotyphla* | **Yes** |
| Hedgehog* | *Erinaceus europaeus* | *Eulipotyphla* | No |
| European mole | *Talpa europaea* | *Eulipotyphla* | No |
| Mountain hare | *Lepus timidus* | *Lagomorpha* | **Yes** |
| European rabbit | *Oryctolagus cuniculus* | *Lagomorpha* | **Yes** |
| Stoat | *Mustela erminea* | *Carnivora* | No |
| Weasel | *Mustela nivalis* | *Carnivora* | No |
| Badger | *Meles meles* | *Carnivora* | **Yes** |
| Otter | *Lutra lutra* | *Carnivora* | No |
| Red fox | *Vulpes vulpes* | *Carnivora* | **Yes** |
| Pine marten | *Martes martes* | *Carnivora* | **Yes** |
| Wildcat* | *Felis silvestris* | *Carnivora* | ? |
| **Peak District** |  |  |  |
| Red deer | *Cervus elaphus* | *Artiodactyla* | **Yes** |
| Roe deer | *Capreolus capreolus* | *Artiodactyla* | No |
| Fallow deer | *Dama dama* | *Artiodactyla* | No |
| Water vole | *Arvicola amphibius* | *Rodentia* | **Yes** |
| Field vole | *Microtus agrestis* | *Rodentia* | **Yes** |
| Wood mouse | *Apodemus sylvaticus* | *Rodentia* | **Yes** |
| Bank vole | *Myodes glareolus* | *Rodentia* | No |
| Brown rat | *Rattus norvegicus* | *Rodentia* | No |
| House mouse | *Mus musculus* | *Rodentia* | No |
| Grey squirrel | *Sciurus carolinensis* | *Rodentia* | **Yes** |
| Harvest mouse* | *Micromys minutus* | *Rodentia* | No |
| Pygmy shrew | *Sorex minutus* | *Eulipotyphla* | **Yes** |
| Water shrew | *Neomys fodiens* | *Eulipotyphla* | **Yes** |
| Common shrew | *Sorex araneus* | *Eulipotyphla* | **Yes** |
| Hedgehog | *Erinaceus europaeus* | *Eulipotyphla* | No |
| European mole | *Talpa europaea* | *Eulipotyphla* | No |
| Mountain hare | *Lepus timidus* | *Lagomorpha* | No |
| Brown hare | *Lepus europaeus* | *Lagomorpha* | No |
| European rabbit | *Oryctolagus cuniculus* | *Lagomorpha* | **Yes** |
| Stoat | *Mustela erminea* | *Carnivora* | No |
| Weasel | *Mustela nivalis* | *Carnivora* | No |
| Badger | *Meles meles* | *Carnivora* | **Yes** |
| Otter | *Lutra lutra* | *Carnivora* | **Yes** |
| Red fox | *Vulpes vulpes* | *Carnivora* | No |
| American mink | *Neovison vison* | *Carnivora* | No |
| Pine marten | *Martes martes* | *Carnivora* | **Yes** |
| Polecat | *Mustela putorius* | *Carnivora* | No |

**Table S2.** Species identified (with a match of ≥0.98 to the reference database) and their associated number of reads after bioinformatic filtering in each site (Assynt A1-A18 and Peak District P1-P3) and in each of three replicates (_1 to _3) for water-based eDNA. The volume of water filtered is indicated for each replicate.

*Additional file: TableS2_Reads_Water.xlsx*

**Table S3.** Species identified (with a match of ≥0.98 to the reference database) and their associated number of reads after bioinformatic filtering in each site (Assynt A1-A18 and Peak District P1-P3) and in each of three replicates (_1 to _3) for sediment-based eDNA. The weight of sediment used for the DNA extraction is indicated for each replicate.

*Additional file: TableS3_Reads_Sediment.xlsx*

**Table S4.** Number of reads obtained after all filtering steps applied to remove non-target MOTUs.

| **WATER** | **Total** |
| --- | --- |
| Total Reads | 13,336,064 |
| After removing reads from the blanks | 10,709,199 |
| After removing non-mammal reads | 10,262,851 |
| After removing human reads | 8,508,564 |
| After removing domestic animals (*Sus, Bos, Equus, Ovis, Canis*) | 5,544,208 |
| MOTUs with minimum identity of 0.98 | 5,414,427 |
| **SEDIMENT** | **Total** |
| Total Reads | 3,309,866 |
| After removing reads from the blanks | 1,684,433 |
| After removing non-mammal reads | 1,543,826 |
| After removing human reads | 649,499 |
| After removing domestic animals (*Sus, Bos, Equus, Ovis, Canis*) | 500,473 |
| MOTUs with minimum identity of 0.98 | 465,997 |

**Table S5**. Mammalian species recorded at seven camera traps in Assynt. Boxes shaded in grey represent sites where each species was recorded.

| **Common name** | **Scientific name** | **Site** | | | | | | |
| --- | --- | --- | --- | --- | --- | --- | --- | --- |
|  |  | A5 | A10 | A11 | A12 | A13 | A14 | A15 |
| Water vole | *Arvicola amphibius* |  |  |  |  |  |  |  |
| Red deer | *Cervus elaphus* |  |  |  |  |  |  |  |
| Field vole | *Microtus agrestis* |  |  |  |  |  |  |  |
| Water shrew | *Neomys fodiens* |  |  |  |  |  |  |  |
| Weasel | *Mustela nivalis* |  |  |  |  |  |  |  |
| Otter | *Lutra lutra* |  |  |  |  |  |  |  |
| Red fox | *Vulpes vulpes* |  |  |  |  |  |  |  |
| Unidentified Shrew | - |  |  |  |  |  |  |  |

**Table S6.** List of tissue samples from mammals used for generating a local reference database using MiFish primers (Miya et al. 2015). All species were tested for amplification using MiMammal-U primers (Ushio et al. 2017) and those highlighted in bold were Sanger-sequenced.

| **Common name** | **Scientific name** | **ID** |
| --- | --- | --- |
| Wood mouse | *Apodemus sylvaticus* | Z.2009.101.1025 |
| Wood mouse | *Apodemus sylvaticus* | Z.2009.101.1149M |
| House mouse | *Mus domesticus* | Z.2009.101.593M |
| House mouse | *Mus domesticus* | Z.2009.101.426 |
| **Field Vole** | ***Microtus agrestis*** | **Z.2009.101.1045** |
| **Field Vole** | ***Microtus agrestis*** | **Z.2009.101.1994M** |
| Bank Vole | *Myodes glareolus* | Z.2009.101.97M |
| Bank Vole | *Myodes glareolus* | Z.2009.101.696M |
| Weasel | *Mustela nivalis* | Z.2009.101.664 |
| Weasel | *Mustela nivalis* | Z.2009.101.363 |
| Yellow-necked mouse | *Apodemus flavicollis* | Z.2009.101.983M |
| Yellow-necked mouse | *Apodemus flavicollis* | Z.2009.101.984M |
| **Water shrew** | ***Neomys fodiens*** | **Z.2009.101.141M** |
| **Water shrew** | ***Neomys fodiens*** | **Z.2009.101.1915M** |
| **Pygmy shrew** | ***Sorex minutus*** | **Z.2009.101.1162M** |
| **Pygmy shrew** | ***Sorex minutus*** | **Z.2009.101.458M** |
| **Common shrew** | ***Sorex araneus*** | **Z.2009.101.611M** |
| **Common shrew** | ***Sorex araneus*** | **Z.2009.101.126M** |
| **Common Vole** | ***Microtus arvalis*** | **Z.2009.101.991** |
| **Common Vole** | ***Microtus arvalis*** | **Z.2009.101.917** |
| Brown Rat | *Rattus norvegicus* | Z.2009.101.931 |
| Brown Rat | *Rattus norvegicus* | Z.2009.101.1026 |
| Grey Squirrel | *Sciurus carolinensis* | 23/24 |
| Grey Squirrel | *Sciurus carolinensis* | 23/10 |
| **Water Vole** | ***Arvicola amphibius*** | **23/15** |
| **Water Vole** | ***Arvicola amphibius*** | **23/17** |
| Edible dormouse | *Glis glis* | 23/16 |
| Edible dormouse | *Glis glis* | 23/35 |
| Brown hare | *Lepus europaeus* | 23/22 |
| Mountain hare | *Lepus timidus* | 23/20 |
| Mountain hare | *Lepus timidus* | 23/1 |
| Hedgehog | *Erinaceus europaeus* | 23/19 |
| Mole | *Talpa europaea* | 23/13 |
| Mole | *Talpa europaea* | 23/14 |
| Red fox | *Vulpes vulpes* | 23/25 |
| Badger | *Meles meles* | 23/12 |
| Badger | *Meles meles* | 23/34 |
| **Otter** | ***Lutra lutra*** | **23/7** |
| **Otter** | ***Lutra lutra*** | **23/33** |
| Polecat | *Mustela putorius* | 23/5 |
| Polecat | *Mustela putorius* | 23/6 |
| Red deer | *Cervus elaphus* | 23/31 |
| Red deer | *Cervus elaphus* | 23/32 |
| Sheep | *Ovis aries* | 23/9 |
| Horse | *Equus caballus* | 24/31 |
| Red Squirrel | *Sciurus vulgaris* | 1/24 |
| Red Squirrel | *Sciurus vulgaris* | 1/31 |
| Pine marten | *Martes martes* | 1/1 |
| Pine marten | *Martes martes* | 1/13 |
| Coypu | *Myocastor coypus* | 62/12 |
| Coypu | *Myocastor coypus* | 22/13 |
| Brown hare | *Lepus europaeus* | 22/7 |
| Stoat | *Mustela erminea* | 22/31 |
| Stoat | *Mustela erminea* | 22/33 |
| Red fox | *Vulpes vulpes* | 21/28 |
| Hedgehog | *Erinaceus europaeus* | 72/32 |
| Sika | *Cervus nippon* | 57/31 |
| Horse | *Equus caballus* | 57/24 |
| Beaver | *Castor fiber* | 63/25 |
| Sheep | *Ovis aries* | 58/31 |
| **American mink** | ***Neovison vison*** | AMX01 |
| **American mink** | ***Neovison vison*** | AMX02 |
| Wildcat | *Felis silvestris* | Z.2015.118.1 |
| Wildcat | *Felis silvestris* | Z.2015.118.2 |

**Table S7**. Maximum number of reads subtracted to control for contamination and/or tag switching for each wild species in each eDNA sampling type (water or sediment) and the type of blank in which the reds were identified (Field, Extraction and PCR). Species indicated by * were not identified as eDNA positive records.

| **Common name** | **Scientific name** | **Blank** | **Reads** |
| --- | --- | --- | --- |
| Red deer | *Cervus elaphus* | Field | 164 |
| Water vole | *Arvicola amphibius* | Extraction | 7479 |
| Field vole | *Microtus agrestis* | Field | 324 |
| Wood mouse | *Apodemus sylvaticus* | None | 0 |
| Brown rat | *Rattus norvegicus* | None | 0 |
| Pygmy shrew | *Sorex minutus* | Field | 1 |
| Water shrew | *Neomys fodiens* | Extraction | 1 |
| Common shrew | *Sorex araneus* | Field | 2 |
| Mountain hare | *Lepus timidus* | Field | 76 |
| European rabbit | *Oryctolagus cuniculus* | Field | 38 |
| Stoat* | *Mustela erminea* | Field | 68 |
| Badger | *Meles meles* | None | 0 |
| Otter | *Lutra lutra* | Extraction | 1 |
| Red fox | *Vulpes vulpes* | None | 0 |
| Pine marten | *Martes martes* | None | 0 |
| Cat | *Felis* spp. | None | 0 |
| American mink* | *Neovison vison* | Extraction | 343 |
| Red squirrel | *Sciurus vulgaris* | Extraction | 1 |
| Grey squirrel | *Sciurus carolinensis* | None | 0 |
| Edible dormouse | *Glis glis* | None | 0 |
| Human 1 | *Homo sapiens* | Field | 547 |
| Human 2 | *Homo sapiens* | Field | 110107 |
| Human 3 | *Homo sapiens* | Field | 1 |
| Cattle | *Bos* spp. | Extraction | 1630 |
| Sheep | *Ovis* spp. | Field | 122 |
| Pig | *Sus scrofa domesticus* | Field | 99 |
| Dog | *Canis lupus familiaris* | Field | 135 |
| Horse | *Equus przewalskii* | None | 0 |

**FIGURES**

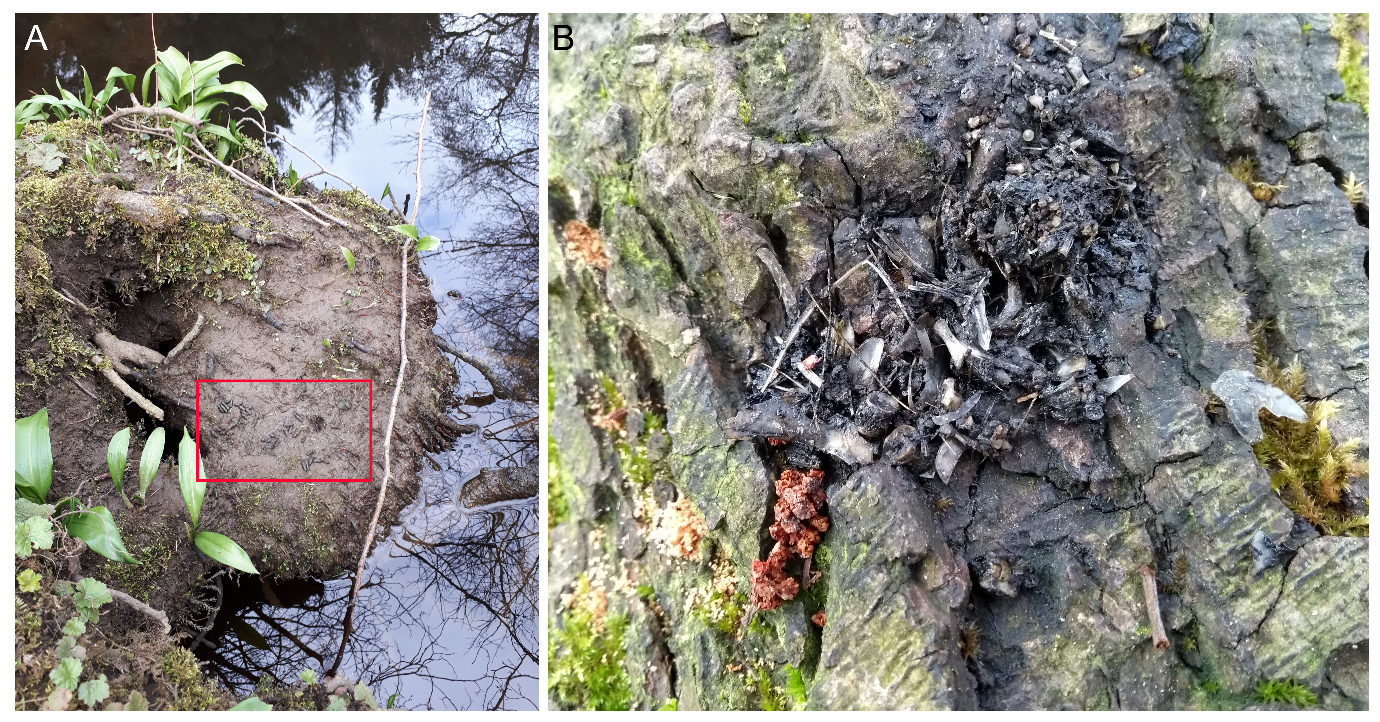

Figure S1. Example of a water vole latrine with faecal pellets, highlighted in the red rectangle in (A), and an otter spraint in (B). Both are from site P1 in the Peak District.

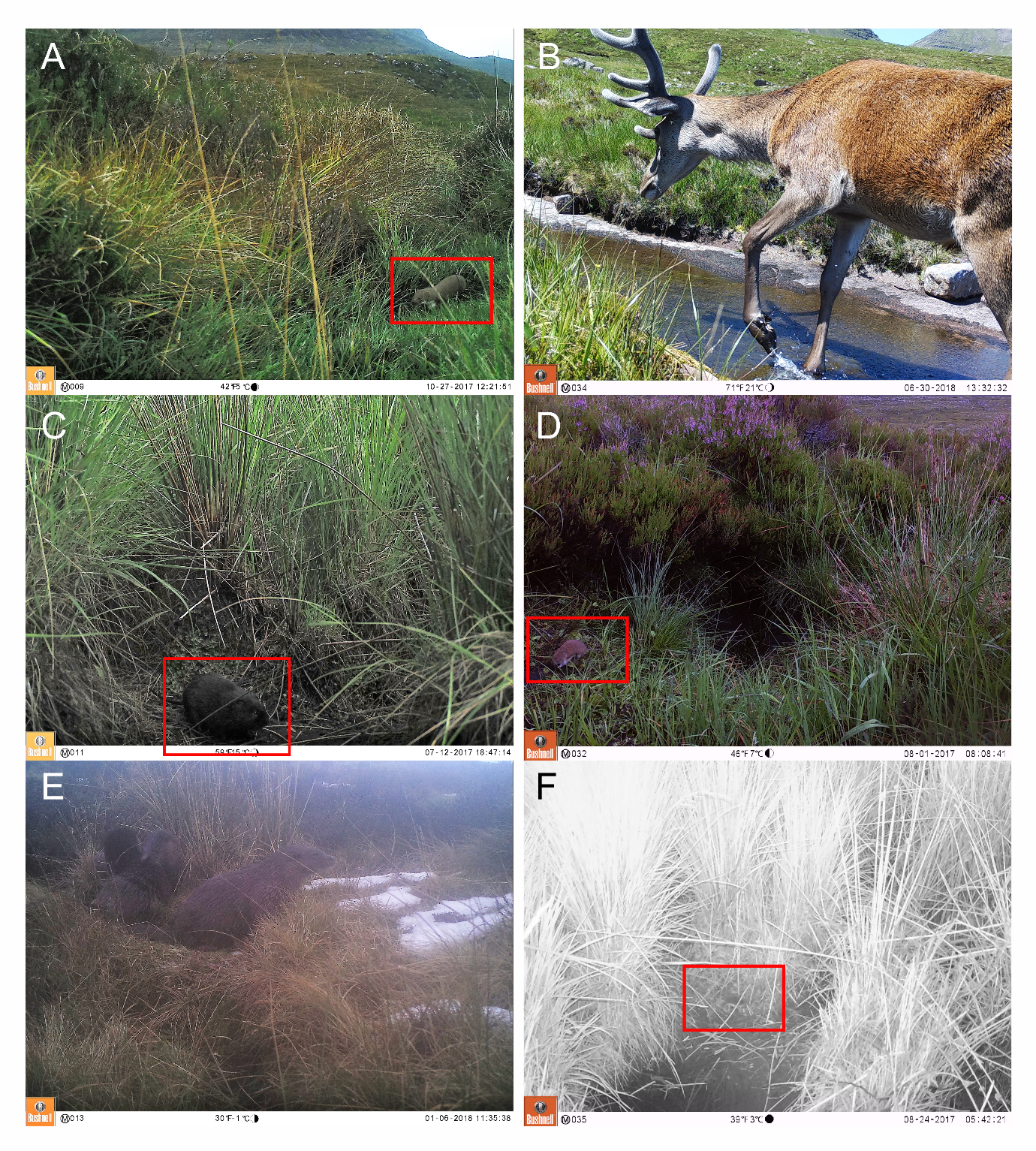

**Figure S2.** Examples of camera trap photographs for six species. Photographs have been manually adjusted to increase visibility of the species. Red boxes are used to highlight where the smaller mammals are positioned within the photograph. A: weasel (*Mustela nivalis*); B: red deer (*Cervus elaphus*); C: water vole (*Arvicola amphibius*); D: field vole (*Microtus agrestis*); E: Eurasian otter (*Lutra lutra*) and F: water shrew (*Neomys fodiens*).

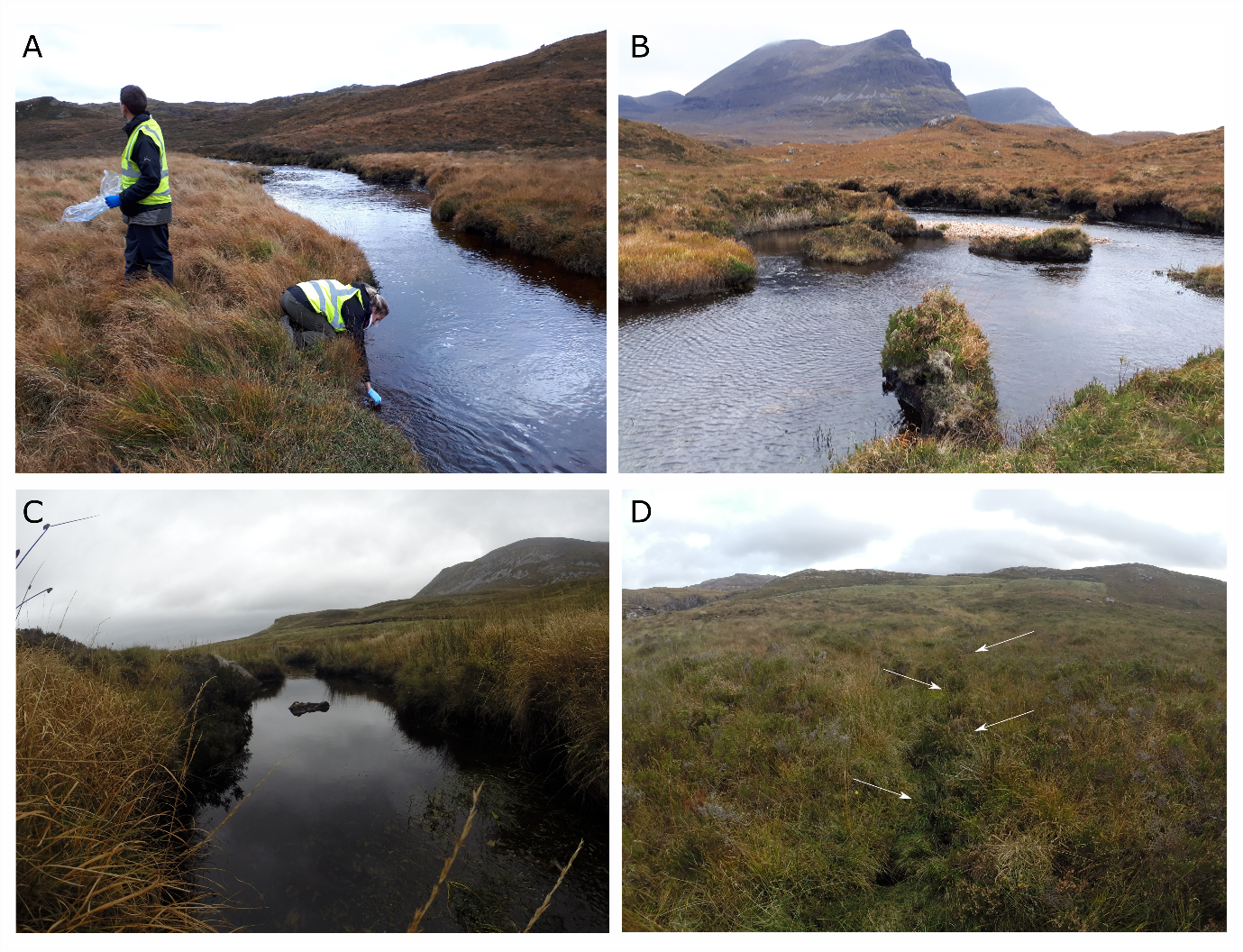

**Figure S3.** Examples of four sampling areas for environmental DNA (eDNA): A = A8; B = A12; C = A16 and D = A11. Sites A8, A11 and A12 returned positive eDNA records for the water vole, site A16 was negative. Sampling at site A11 was conducted in a narrow stream that is not visible here but is indicated by the white arrows (D). Sampling methodology for eDNA is indicated in (A), where sampling was conducted along the edge of the river/stream for both water and sediment samples.

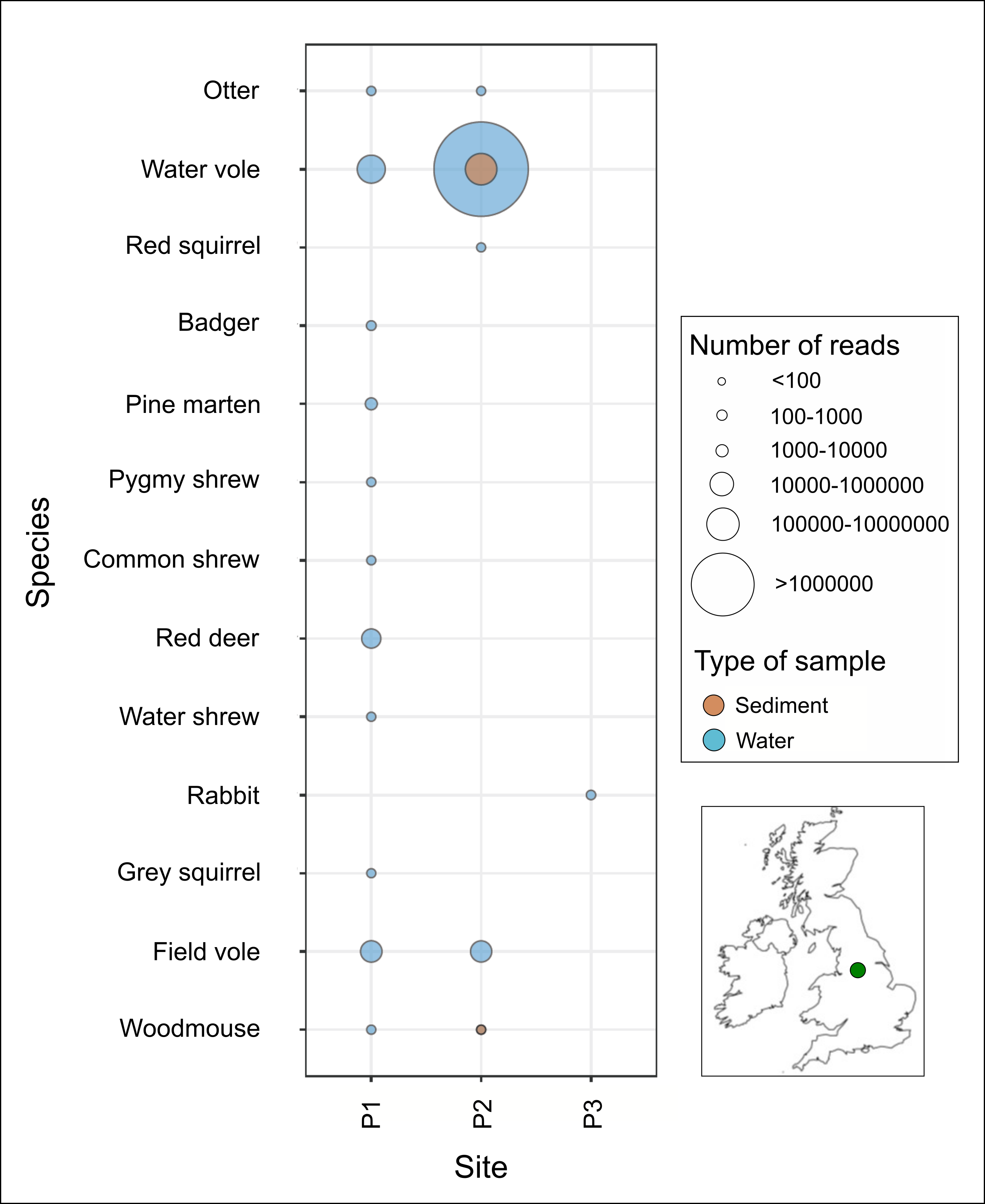

**Figure S4:** A bubble graph representing presence-absence and categorical values of the number of reads retained (after bioinformatic filtering) for eDNA (water in blue and sediment in orange) from each wild mammal identified in each site (P1-P3) in the Peak District National Park. The location of the Peak District is indicated in the inset map but the actual sampling sites can not be disclosed due to conservation and persecution concerns around certain protected species.
